# A Scalable 3D High-Content Imaging Protocol for Measuring a Drug Induced DNA Damage Response Using Immunofluorescent Sub-nuclear γH2AX Spots in Patient Derived Ovarian Cancer Organoids

**DOI:** 10.1101/2022.09.15.508096

**Authors:** Hakan Keles, Christopher A. Schofield, Helena Rannikmae, Erin Elizabeth Edwards, Lisa Mohamet

## Abstract

The high morbidity rate of ovarian cancer has remained unchanged during the past four decades, partly due to lack of understanding of disease mechanisms and difficulties in developing new targeted therapies. Defective DNA damage detection and repair is one of the hallmarks of cancer cells and is a defining characteristic of ovarian cancer. Most in vitro studies to date, involve viability measurements at scale using relevant cancer cell lines, however, the translation to clinic is often lacking. The use of patient derived organoids is closing that translational gap yet the 3D nature of organoid cultures present challenges for assay measurements beyond viability measurements. In particular, high-content imaging has the potential for screening at scale providing a better understanding of mechanism of action of drugs or genetic perturbagens. In this study we report a semi-automated and scalable immunofluorescence imaging assay utilising the development of a 384-well plate based subnuclear staining and clearing protocol and optimisation of 3D confocal image analysis for studying DNA damage dose response in human ovarian cancer organoids. The assay was validated in four organoid models and demonstrated a predictable response to Etoposide drug treatment with lowest efficacy observed in the clinically most resistant model. This imaging and analysis method can be applied to other 3D organoid and spheroid models for use in high content screening.

## INTRODUCTION

Ovarian cancer is known to be a heterogenous disease that consists of multiple distinct malignancies that share a common anatomical site. Of these, the type II high-grade serous (HGS) subtype dominates in the clinical setting and is responsible for over 70% of all cases ^1 2 3 4^. The high morbidity rate and the finding that the 5-year overall survival from ovarian cancer has remained virtually unchanged since about the 1980s, is due to factors including late detection of disease, lack of understanding of disease mechanisms and difficulties in developing new targeted therapies ^1, 2^. Currently the primary route of therapy is surgery and chemotherapy, with targeted therapies mainly utilised in recurrent disease, both in chemotherapy sensitive and resistant cases ^1^.

Effective targeted therapies rely on knowledge of the molecular features of the disease. Defective DNA damage detection and repair (DDR) is one of the hallmarks of cancer cells and is a defining characteristic of HGS ovarian cancer ^5^. Defects in the homologous recombination repair (HRR) pathway have been found in more than 50% of patients with HGS ovarian cancer, with BRCA1 and BRCA2 playing particularly pronounced roles ^6 7 8^. This feature of HGS ovarian cancers has been utilised for targeted therapies as part of the concept of synthetic lethality, where functional loss of individual genes is tolerated in isolation but not in combination ^5^. Inhibition of poly (ADP-ribose) polymerase proteins (PARPs) - nuclear enzymes activated by DNA damage and integral to DNA repair - is one such example, where ovarian cancers with defects in HRR pathway display synthetic lethality in the context of PARP inhibition ^9^.

Preclinical human models of ovarian cancer include cancer cell lines, patient derived xenografts (PDX) and organoids. Cancer cell lines serve well in terms of high scalability and low cost and (still) contribute to the discovery of novel targets ^10^. However, they lack translatability to the patient due to numerous factors ranging from their initial derivation, clonality, adaptation to *in vitro* 2D growth, genetic drift, and cross contamination amongst other factors ^11^. PDX offer increased genetic and histological stability ^12^ and have been used successfully in pre-clinical research ^13 14 15^ however, their use is ethically challenged, labour intensive, and lacks the scalability needed for early drug screens.

Patient derived organoids demonstrate stable phenotype expression in a 3D culture platform that have been shown to recapitulate tumour heterogeneity ^2 3^. They are scalable and recapitulate the patient’s response to chemotherapeutics in vitro ^4, 16, 17^. The origin of the majority of HGS is widely acknowledged to be from the epithelium of the fallopian tube ^1^ which is highly relevant for organoid culture as these are derived from epithelium. Organoids are in use in pre-clinical screening assays ^17^ however, such assays have typically been limited to cell viability at such scale. Although viability assays provide a robust and reliable readout for oncology, additional information regarding the cellular machinery or pathways involved in cell killing could increase our understanding of the disease and aid in the identification of novel molecular targets for potential therapies. Due to the 3D and heterogenous architecture of organoids and the frequent use of basement membrane extracts for their culture, the options of cellular interrogation at scale in a multi-well format are limited. Automated confocal high-content imaging is one option that can be used to overcome this challenge in organoid culture system, however, requires thoughtful consideration insofar as how to amend conventional 2D high content imaging workflows to accommodate this challenging 3D context.

High content screening (HCS) is an integral part of drug discovery platforms, and it is widely used to identify and validate compounds by classification of drug induced phenotypes at high throughput. A HCS method for the quantification of DNA damage utilising γH2AX labelled double strand DNA break foci as a marker was reported in cancer cells with a large-scale chemical library in 2D imaging format ^18^. However, with increasing evidence of improved clinical relevance in the assays utilising primary human tissue in the form of organoids compared to that of 2D cell lines, there is growing need for scalable methods for high content confocal imaging screens in 3D. The multivariate nature of the phenotypic imaging data is crucial for cross validating established assays such as viability or similar in patient derived 3D cellular models like organoids ^19^.

However, utilisation of 3D cell culture in HCS is still challenging primarily due to difficulties in scalability to 384-well plates which are crucial for leveraging the full benefits in terms of assay robustness and throughput that automated screening setups provide ^20^ as well as for minimising excessive use of difficult to scale organoid source cultures. As such, liquid handling protocols need to be tuned not only to handle often delicate complex cultures, but also to do so in low volumes. With respect to imaging, the combination of organoid size, mismatched refractive indices of cell and organoid sized structures, and encapsulation in hydrogels such as Matrigel present unique challenges, that require careful optimisation of labelling protocols as well as optical clearing to ensure sufficient label and laser penetration.

Recent publications demonstrate some advances in the industrialisation of 3D complex cell based assays, for example, monitoring kidney organoid differentiation from human pluripotent stem cells ^21^, monitoring growth in response to Tankyrase inhibitors ^22^ and quantifying compound related cytoskeletal and nuclear phenotypic changes in colorectal organoids ^23^. However, to the best of our knowledge, there is still a need for a reproducible multi-well plate based immunofluorescence imaging protocol for compound induced 3D DNA damage response imaging at scale in ovarian cancer organoids.

Utilisation of organoids in a 3D image based HTS in 384-well plates requires optimisation of cell culture parameters, compound treatment regimen, assay time course and endpoint staining conditions. Furthermore, such protocol optimisation also requires robust and scalable assays that are amenable to multiple liquid handling devices and data pipelines in place to enable optimised image acquisition parameters and image analysis workflows for a scalable 3D immunofluorescence method.

Several organoid or spheroid fixing, and staining protocols are reported to improve signal to noise ratio (SNR) in 3D fluorescence microscopy images with increased laser penetration depth facilitating quantification of fluorescence signal in 3D multicellular structures ^24^. We have modified and adapted a method reported for cells on slides or dishes for high resolution confocal laser scanning microscopy. 384-well assay plates were utilised for automated confocal microscopy with automated liquid handling to enable the scale required for screening.

Here, we report the development of a scalable 3D imaging assay using patient derived HGS ovarian cancer organoids. We quantified the compound-initiated 3D DNA damage response using a subnuclear immunofluorescence assay using a γH2AX antibody. 69 intensity, morphology and texture related parameters were then calculated for organoids, organoid nuclei and subnuclear γH2AX spots, in batch mode through an automated analysis script.

DNA damage response to Etoposide was used to validate the high-content imaging method by calculating Z-prime scores using, DMSO and 30 μM Etoposide, as negative and positive controls, respectively. The largest effect size for DNA damage response was seen for the parameter ‘Total nuclear γH2AX spot area- mean per well’ for the organoid lines studied. In addition, phenotypic parameters such as those related to γH2AX spot texture, organoid nuclear size and nuclear roundness were seen to be dependent on compound concentration thus demonstrating the value of phenotypic high-content imaging in preclinical drug discovery as a multivariate screening tool. Furthermore, one of the organoid lines did not respond to Etoposide confirming its resistant nature.

## MATERIALS AND METHODS

### Patient-Derived Ovarian Cancer Organoid Culture

Cryopreserved ovarian tumour organoids were obtained from Hubrecht Organoid Technology (HUB; Netherlands). Figure 1 shows brightfield microscopic images of the models in culture, they show a variety of compact morphologies and were derived from HGS carcinomas from different patients, sites, and stages of disease (Table 1). The human biological samples were sourced ethically, and their research use was in accord with the terms of the informed consents under an IRB/EC approved protocol.

**Figure 1.**
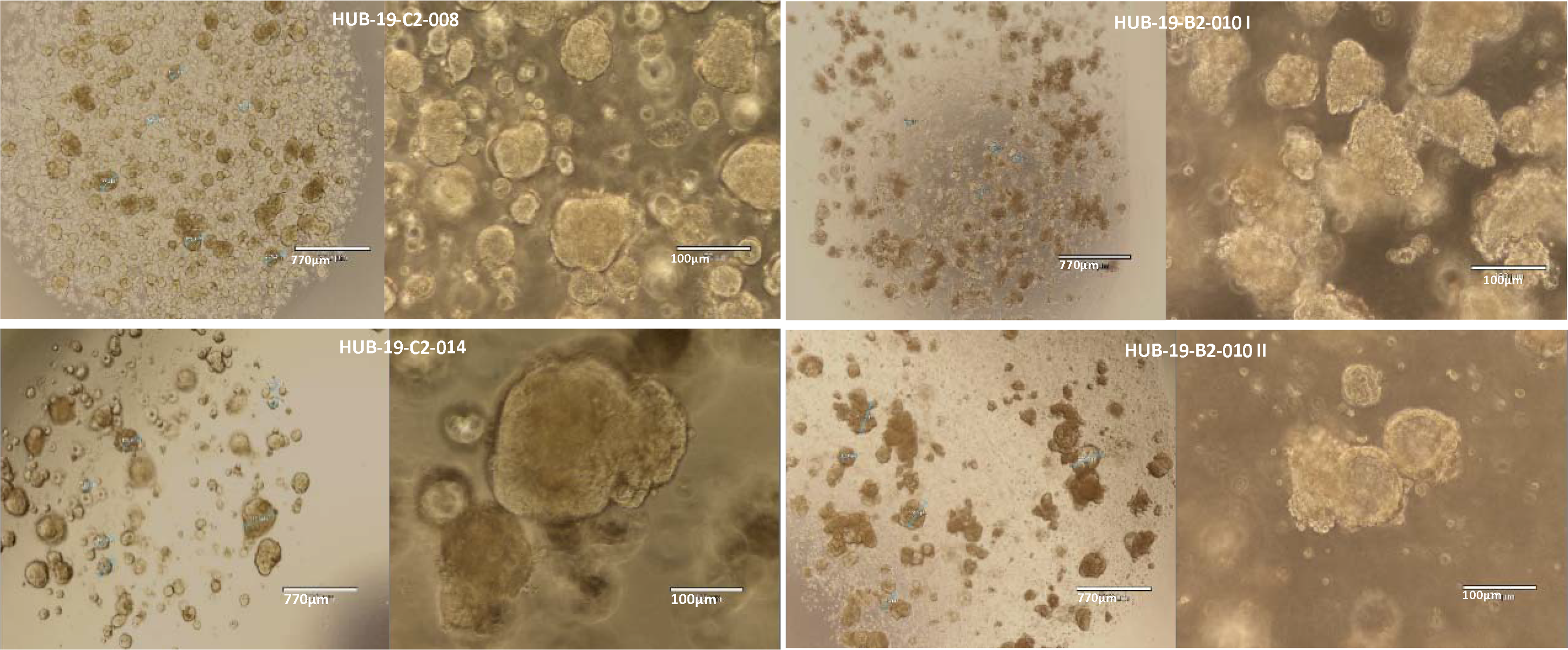
4x magnification (left) and10x magnification (right) brightfield microscopic images of the 4 ovarian cancer organoid lines in Matrigel droplets after 7 days culture.

**Table 1.** List of organoid lines used.

| HUB code | Abbreviation | Patient | Disease | Donor (sex, age, site of collection) | Organoid morphology |
| --- | --- | --- | --- | --- | --- |
| HUB-19-C2-008 | HUB008 | 1 | HGS carcinoma, original tumour | F, 70, omentum | Compact |
| HUB-19-C2-014 | HUB014 | 1 | HGS carcinoma, recurrent tumour | F, 71, ascites | Compact |
| HUB-19-B2-010 I | HUB010 | 2 | HGS carcinoma | F, 67, adnex right | Compact |
| HUB-19-B2-010 II | HUB010 II | 2 | HGS carcinoma | F, 67, adnex left | Compact |

In Figure 1, all organoids have the expected compact morphology with a wide range in size between approximately 30 μm and 300 μm in diameter depending on the extent of shearing used during sub-culture, subsequent growth rate and aggregation. Patient 2 organoids (HUB010, HUB010II) were less compact than Patient 1 (HUB008, HUB014) making them easier to shear into smaller fragments. The concentration of the organoids is variable but controlled and adjusted as needed for optimum growth by the split ratio applied at the time of sub-culture.

Cultures were maintained and expanded in 10 μl droplets of 80% Matrigel (Corning, Cat# 356231) in 6-well suspension plates (Greiner, Cat# 657185) and cultured in ovarian cancer (OC) medium without Wnt as previously described ^3^ except recombinant human R-spondin 3 (R&D Systems, Cat# 3500-RS-025/CF) was used at 25 μg and recombinant human Noggin (Peprotech, Cat# 120-10C) was used at 20 μg for 100 ml of complete medium.

Ovarian cancer organoids were subcultured with a ratio of 1:3 or 1:2, dependant on the donor line, every 7 days by mechanical shearing using a filtered P1000 pipette tip with a non-filtered P10 pipette tip on the end. Large and dense organoids were sheared as before but in pre-warmed (37 °C) 50% TrypLE (Gibco, Cat# 12604013) + 50% ADF+++ (Advanced DMEM/F-12 (Thermo Fisher, Cat# 12634010), PenStrep (Thermo Fisher, Cat# 15140122) and GlutaMax (Gibco, Cat# 35050061)) buffer. Sheared organoid pellets were washed with ADF+++, spun 450 *g* at room temperature for 5 minutes and any old remaining Matrigel removed before resuspending in fresh 80% Matrigel and re-plating. The plates containing freshly plated droplets were inverted and incubated at 37°C, 5% CO_2_ in air for 30 minutes until Matrigel had polymerized and then turned the right way up before adding 2 ml OC medium and re-incubating. OC medium was replaced every 2-3 days.

### Organoid Expansion for HCS

Organoids were expanded as described above to generate 1 × 10^6^ organoids of 20 μm – 70 μm diameter needed for the assay. This equated to seven confluent 6-well suspension plates per donor line (total of 42 wells with 18 × 10 μl droplets per well). Two days before seeding for the assay, the organoids were subcultured and sheared with a ratio of 1:1 into 10 μl droplets of 50% Matrigel.

1. Collect culture plates of ovarian organoids from the incubator and check confluency, organoid size, growth, and general health of the cultures by light microscopy at 4x to 10x magnification.
2. Add 20 ml of Dispase stock (100 mg/ml) to each well (already containing 2 ml culture medium) and return to incubation (37 °C, 5% CO_2_, in air) for 60 minutes.
3. Prepare sufficient OC medium containing 5% Matrigel and store on ice for use in step 16.
4. Prepare the Multidrop combi instrument, use ice cold DPBS to chill the tubing just prior to plating in step 17.
5. Pre-wet sufficient 70 μm cell strainers (pluriSelect, Cat# 43-50070-01) with 1 ml D-BSA on each side of the filter. One filter for up to 4 full 6-well plates.
6. Collect cultures after 60 minutes incubation and check Matrigel domes have dissolved releasing the organoids into the culture supernatant.
7. Aspirate the supernatant containing organoids from all wells and pass through the 70 μm cell strainer into a 50 ml centrifuge tube.
8. Wash each culture well with 2 ml D-BSA and use to wash through the cell strainer keeping the final volume less than 50 ml.
9. Pre-wet sufficient number of 20 μm cell strainers (pluriSelect, Cat# 43-50070-01) with 1 ml D-BSA on each side of the filter. One filter for up to 4 full 6-well plates.
10. Pass the <70 μm flow through over the 20 μm cell strainers, wash the strainers with 10 ml with ADF+++.
11. Collect the 20 μm – 70 μm size organoids by back-washing them from the top surface of the 20 μm cell strainer into a separate 50 ml centrifuge tube with ADF+++.
12. Spin these organoids into a single pellet at 450 *g* for 5 minutes at room temperature.
13. Discard the supernatant and re-suspend the pellet in 0.5 ml to 2 ml ice cold OC media containing 5% Matrigel depending on the expected count. Keep the tube on ice during the count.
14. Gently, but thoroughly, mix the organoid suspension then take a 20 μl sample and dilute with 180 μl ADF+++.
15. Count the number of organoids using the CytoSMART counter (Corning) as per the manufacturer’s instructions.
16. Calculate the total number of organoids and dilute to a concentration of 20,000 organoids per ml using ice cold OC media containing 5% Matrigel.
17. Chill the multidrop combi tubing by flushing through with ice cold DPBS.
18. Mix the organoid suspension gently but thoroughly before priming the multidrop combi tubing and adding 20 μl suspension (400 organoids) to all wells of the 384-well assay plates (PerkinElmer cell carrier Ultra sterile and ultra-low-attachment-coated plates, Cat# 6057802).
19. Prior to incubation, replace the standard plate lids with environmental microclime lids (Labcyte, Cat# LLS-0310) following the manufacturer’s instructions.
20. Prepare 5 replicate plates containing the test conditions.
21. Incubate the plates in a humidified atmosphere of 5% CO_2_ at 37 °C for 1hour before addition of compound.

### Etoposide Treatment

Etoposide (DNA Topoisomerase II) was prepared and stored frozen as a concentrated DMSO (DIMETHYL SULFOXIDE) stock prior to dilution and addition to the organoids. The compound was serially diluted in 2- fold steps from the highest test concentration (specified in Table 2) and assayed over 7 concentrations with a final assay v/v DMSO concentration of 0.5%.

**Table 2.** Etoposide dilution range.

| Compound Number | CAS No. | Compound Name | Highest final concentration | Lowest final concentration | Dilution Series |
| --- | --- | --- | --- | --- | --- |
| GR87698X | 33419-42-0 | Etoposide | 30 µM | 470 nM | 1 in 2 |

Compound plate preparation for addition to organoids was performed using the following method.

1. Thaw the compound plate at room temperature for 30 minutes.
2. During this time prepare sufficient OC media containing 5% Matrigel and store on ice for use in step 4.
3. After 30 minutes, centrifuge compound plates at 50 *g* for 1 minute at room temperature.
4. Dilute the compound plate to 2x final assay concentration with OC media containing 5% Matrigel using the BRAVO (Agilent) liquid handler.
5. Centrifuge diluted compound plates at 50 *g* for 1 minute at room temperature.
6. Mix the well contents and transfer 20 μl of diluted compound to the organoid plate using the BRAVO liquid handler. Total volume per well will be 40 μl with compounds at their final assay concentration.
7. Image the plates with a 5x objective in brightfield transmission mode using PerkinElmer Opera Phenix high content screening system.
8. Incubate the plates as described previously for 48 hours.

### CellTitre-Glo 3D Cell Viability Assay

Organoid viability was measured in one of the 5 replicate plates using the CellTitre-Glo 3D Cell Viability kit (Promega, Cat#G9683) using the following method.

1. Image the whole 384-well plate using PerkinElmer Opera Phenix high content screening system in brightfield transmission mode with the 5X lens to check well contents.
2. Thaw 40 ml CellTiter-Glo 3D and equilibrate to room temperature.
3. Dispense and mix 30 μl of CellTiter-Glo reagent in each well using the BRAVO liquid handler.
4. Incubate in the dark at room temperature for 30 minutes by wrapping the plate in foil.
5. After 30 minutes, mix and transfer 10 μl to a solid white low volume assay plate (Greiner, Cat# 784080) using the BRAVO liquid handler.
6. Measure the luminescence from the whole 384-well plate using a PHERAstar® FSX plate reader (BMG Labtech).
9. Calculate percent viability of treated wells using DMSO as the high control.

### Organoid Staining and Clearing for Imaging

Organoids were fixed after 48 hours of incubation with Etoposide in media following the steps below, using consumables, reagents and devices listed. A visual summary of this workflow can be seen in Supplementary Figure S1.

#### Reagents

All reagents were prepared fresh and filtered on the day using consumables listed in Supplementary Table S1.

1. OWB (organoid washing buffer); 500 ml DPBS + 1 g BSA + 0.5 ml Triton X-100.
2. PBT (DPBS with Tween); 500 ml + DPBS + 0.5 ml Tween.
3. DPBS-BSA; 500 ml DPBS + 1 g BSA.
4. Clearing Stock1; (50% (v/v) MeOH + 25% (v/v) DPBS + 25% (v/v) H_2_O).
5. Clearing Stock2; (80% (v/v) MeOH + 20% (v/v) H_2_O).
6. Clearing Stock3; (100% MeOH).

#### Equipment

1. Corning® LSE™ Digital Microplate Shaker (kept and used in a 4°C cold-room).
2. BRAVO automated liquid handler (Agilent Technologies).
3. Viaflo384 liquid handler (Integra Biosciences).

Plates were fixed, stained and cleared after 48 hours of compound incubation as below.

1. Image the plates with a 5x objective in brightfield transmission mode using PerkinElmer Opera Phenix high content screening system.
2. Visually inspect the plates for viability, size, and distribution of organoids in each well (40 μl media) before further processing.
3. For fixing, spin plates at 200 rpm for 10 seconds and then place on the BRAVO liquid handler for 5 minutes to settle.
4. Wash BRAVO tips with 40 μl DPBS-BSA once, dispense 40 μl, 4% PFA ready solution into each well.
5. Incubate *in situ* for 15 minutes.
6. Aspirate 40 μl from each well and discard.
7. Add another 40 μl 4% PFA to each well as before.
8. Incubate *in situ* for 15 minutes.
9. Aspirate 40 μl from each well and discard.
10. Wash wells with 40 μl PBT twice in a fume hood using a BRAVO.
11. Add 40 μl PBT and if required store the plates at 4 °C (for up to 72 hours).
12. Aspirate 40 μl PBT, add 40 μl OWB.
13. Incubate *in situ* for 15 minutes.
14. Aspirate and replace with another 40 μl of OWB.
15. Incubate on a shaker (70 rpm, no tilt) at 4 °C in fridge for 60 minutes.
16. Spin the plates at 200 rpm for 10 seconds.
17. Incubate for 15 minutes at room temperature and aspirate 40 μL OWB.
18. Add 40 μl primary antibody solution (prepared in OWB) i.e., γH2AX (2:1000) and incubate overnight on a shaker (70 rpm, no tilt) at 4 °C in dark fridge.
19. Spin the plates at 200 rpm for 10 seconds.
20. Incubate on BRAVO at least 15 minutes then aspirate and discard 40 μl primary antibody solution.
21. Add 40 μl OWB and incubate *in situ* for 15 minutes, then aspirate and discard.
22. Add 40 μl OWB and incubate on a shaker (70 rpm, no tilt) at 4 °C in fridge for 1 hour.
23. Place the plates on the BRAVO for at least 15 minutes then aspirate 40 μl OWB.
24. Add 40 μl of the secondary antibody i.e., Alexa 647 (2:1000) solution and dapi (2:1000) stain and incubate overnight on a shaker (70 rpm, no tilt) at 4 °C in dark fridge.
25. Spin the plates at 200 rpm for 10 seconds.
26. Incubate on BRAVO at least 15 minutes then aspirate and discard 40 μl secondary antibody solution.
27. Add 40 μl OWB and incubate *in situ* for 15 minutes, then aspirate and discard.
28. Add 40 μl OWB and incubate on a shaker (70 rpm, no tilt) at 4 °C in fridge for 1 hour.
29. Place the plates on the BRAVO for at least 15 minutes then aspirate 40 μl OWB.
30. Add 40 μl OWB and incubate *in situ* for 15 minutes, then aspirate and discard.
31. Add 40 μl stock1 using a Viaflo384 in a fume hood. Incubate at room temperature (RT) 15 minutes.
32. Aspirate 40 μl stock1, add 40 μl stock 2 using a Viaflo384 in a fume hood and incubate at RT for 15 minutes.
33. Aspirate 40 μl stock2, add 40 μl stock 3 using a Viaflo384 in a fume hood and incubate at RT 15 minutes.
34. Aspirate 40 μl stock3, add 40 μl clearing reagent and incubate the plate at room temperature for 15 minutes.
35. Record a 5x brightfield image of the plate for visual checks of losses after protocol
36. Seal the plate and move to imaging and analysis.

### Automated 3D High-Content Imaging

A PerkinElmer Opera Phenix high content screening system coupled with a collaborative robot designed for pharmaceutical screening (the plate:handler™ FLEX, PerkinElmer) was used to image the 384-well plates in spinning disk confocal mode. A 20x water immersion objective (NA=1.0) was used to collect 16 fields of view from each well covering ~70% of the total well area, utilising two fluorescence channels comprising nuclei (excitation: 405 nm, emission: 435 nm-480 nm) and γH2AX spots (excitation: 640 nm, emission: 650 nm-760 nm). A Z-stack of 14 focal planes were collected from each field of view with 10 μm Z-steps. Two sCMOS cameras of the system were employed with pixel binning of 2. Each camera acquired one fluorescence channel at a time, resulting in 3 hours scan time per full 384-well plate (~270 GB file size). All fluorescence imaging data was collected, visualized, and analysed using Perkin Elmer Harmony (version 4.9.2137.273, Revision: 147881, Acapella version: 5.0.1.124082) software installed on the imaging instrument and analysis computers. Raw image files were annotated within the software via definition of links for electronic experiment records and plate maps with compound names, compound concentration and cell donor codes ensuring meta data was FAIR (Findable, Accessible, Interpretable, and Reusable).

### Image Analysis

Images were analysed using a second copy of the Harmony software package installed on a local PC (HP Z8 G4 workstation, Windows 10 (64-bit), dual intel Xeon6248R 3.0 GHz CPU, Nvidia QDR 16 GB TRX 5000 Graphics, HP Z turbo 4×1 TB SSD, 192 GB DDR4 RAM) connected to the imager PC (DELL workstation, similar specification) via 10 Gigabit ethernet. Analysis of a full 384-well plate took ~3 hours which at the time of this study was relatively shorter than running the analysis on a cluster using PerkinElmer’s Columbus 2.9 platform, mainly due to low speed of file mounting from Harmony to Columbus. Raw images were flat field corrected using the basic correction algorithm available in the Harmony image analysis package. A maximum intensity projection of the collected Z-stacks was obtained for each field of view. A set of representative images showing nuclei in blue and γH2AX in red, from both negative (DMSO) and positive (30 μM Etoposide) control wells can be seen in Figure 2. In Figure 2 (D), DNA damage can be seen expressed as various size spots and larger areas in some nuclei or co-localised with up to 100% of the individual nuclei. This variation in γH2AX expression is expected compared to other more often communicated DNA damage inducing methods such as irradiation. Therefore the spot detection algorithm that is able to distinguish such spots with different morphology and intensity, needed to be implemented by editing avaiable options in the Harmony software.

**Figure 2.**
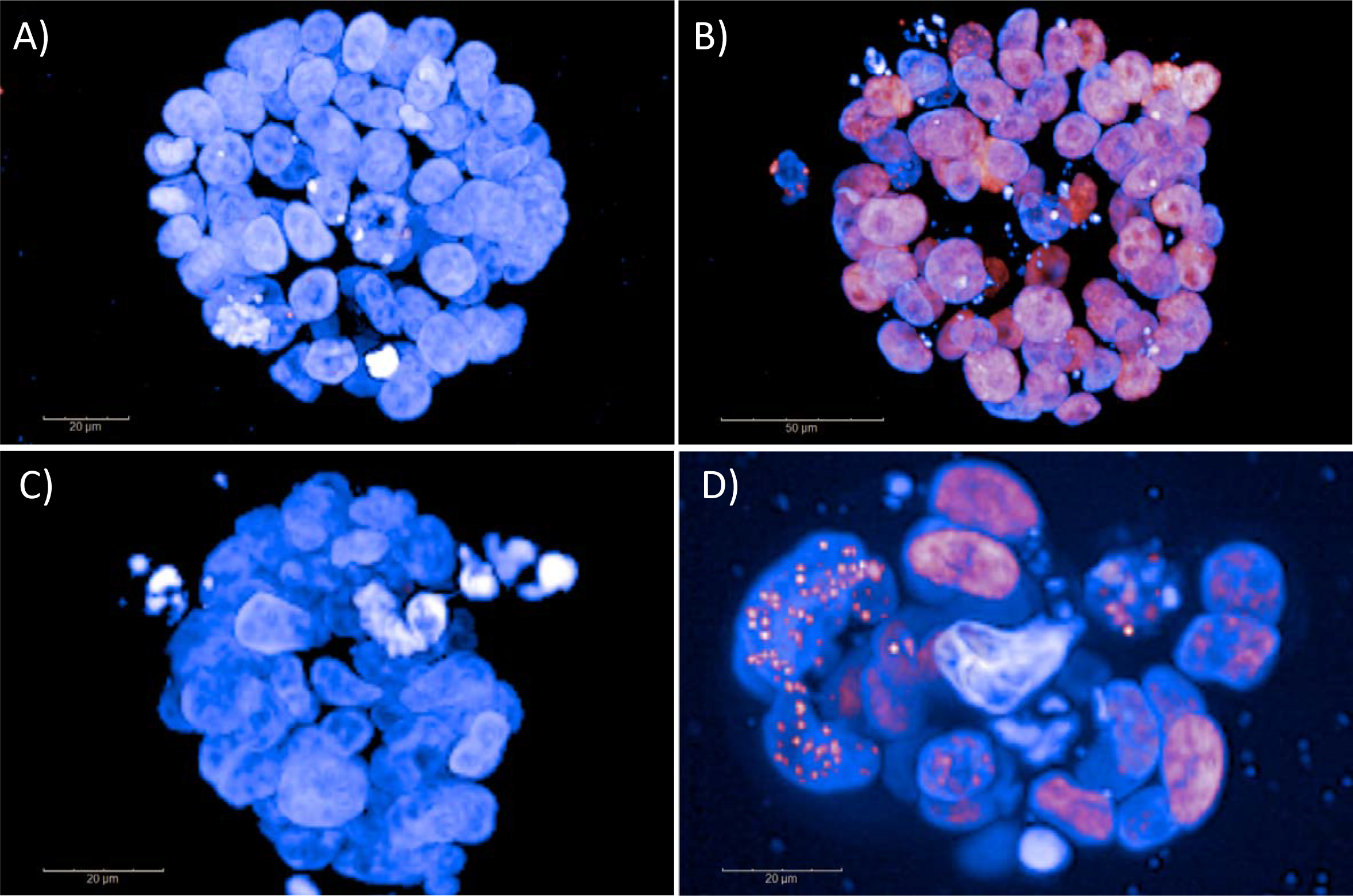
Maximum intensity projected control (A and C) and 30 μM Etoposide treated (B and D) images of the HUB008 organoid line.

Figure 3 outlines the image analysis steps. Briefly, the nuclear channel in the projected images was used to detect organoids larger than 800 μm^2^ and those touching edges of the images were removed from analysis. The rest of the organoids were further filtered based on roundness (>0.01), average DAPI intensity (>500) and width to length ratio (>0.03) in order to remove debris from the wells (dust, fibres, deplated organoids etc.). Selected organoids were then used for nuclear segmentation using nuclei detection method ‘M’ and spots were detected within individual nuclei using the spot detection method ‘D’, which was seen to be distinguishing the variable shapes and spot sizes, avaiable in the Harmony analysis software. Intensity, morphology and texture related parameters were then calculated for organoids, organoid nuclei and subnuclear γH2AX spots utilsing the automated analysis pipeline that was running in parallel to imaging.

**Figure 3.**
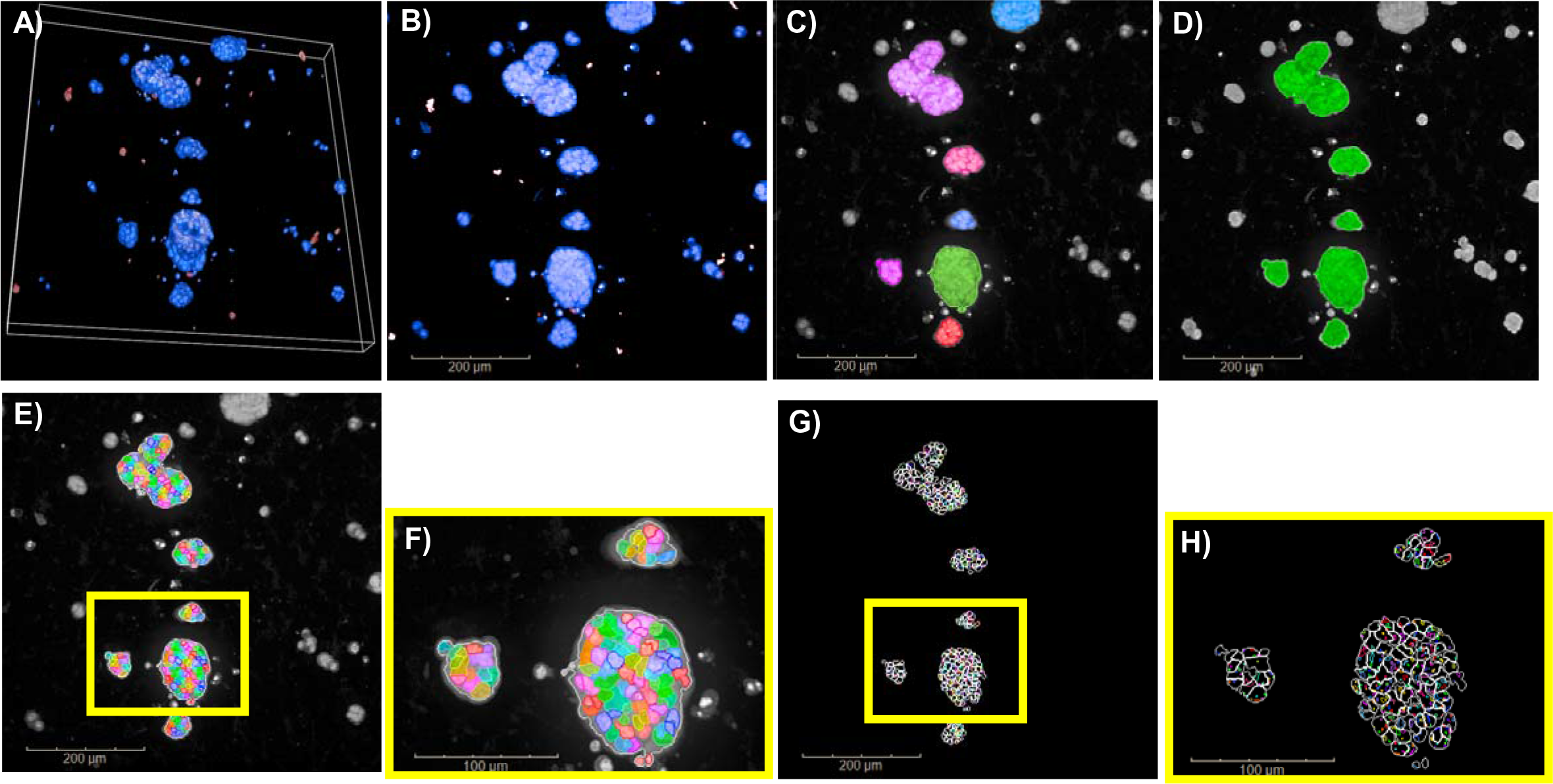
3D Visualisation (A) and analysis workflow as applied in one field of view of a maximum intensity projected Z-stack (B). Detected organoids (C) are filtered (D) and segmented (E) and (F). γH2AX spots are then detected in individual nuclei G) and (H).

### Plate and Well Quality Control

Image analysis result files, comprising a list of intensity and morphology features for each plate with annotations readily added in Harmony software, were exported in text format and imported into High Content Profiler v2.0.0 application (PerkinElmer) in Tibco® Spotfire® package v11.4 to be used for secondary analysis and data visualization. All plates were analyzed at once, using default settings in the High Content Profiler application mainly in that an inter-plate normalization was applied using the ‘median method’ and feature selection was done comparing Z-prime scores ^25^.

All of the calculated features were used individually to calculate a Z-prime score indicating effect size in each plate using the difference between negative (DMSO) and positive (30 μM Etoposide) controls. For High content screening assays Z-prime scores above 0 are considered sufficient, depending on the complexity of the assay ^25^. Total nuclear γH2AX spot area (mean per well) was found to be the highest Z-prime scoring parameter across all responding organoid lines. A list of parameters can be seen in Supplementary Table S2 (B). All dose related plots and donor comparisons were then generated based on the highest Z-prime scoring parameter ‘Total nuclear γH2AX spot area- mean per well’.

Each organoid line was imaged in 4 replicate plates comprising 32 x DMSO, 32 x Etoposide (30 μM) replicates, and 8 × 7-point Etoposide dose response curves. Raw Etoposide 7-point dose response data plotted for the highest Z-prime scoring parameter (across all lines) is shown in Supplementary Figure S4. A dose response curve can readily be seen for HUB008 (Patient 1 original tumour), HUB010 (Patient 2 adnex right) and HUB010 II (Patient 2 adnex left). No remarkable response was observed in the HUB014 line (Patient 1 recurrent tumour). The raw data for HUB010 II shows the largest amount of variability overall, including outliers.

Supplementary Table S2 (A) shows the list of Z-prime and SNR scores calculated for each plate for the 3 organoid lines that showed a dose response in raw data shown in Supplementary Figure S2. A quality control threshold was applied such that outlier wells that showed edge effects, including less than 5 organoids, were removed from analysis to cut false signal contribution to the rest of the wells (on average ~50 organoids per well was detected). An average signal to noise ratio of 4.2 was achieved across all the plates which is acceptable considering the 3D nature of the assay, since projecting images for maximum intensity per binned pixel across all Z-planes results in increased signal but also increases noise for some binned pixels if they do not contain organoids in that particular field of view. Selective imaging (i.e., pre-scan with low magnification lens (5x or 10x) to locate and re-scan with high magnification lens (20x water) to resolve individual organoids or volumetric analysis can be implemented to improve SNR however these options come at the expense of computational power and time which at the time of our study risked the practicality and scalability of the reported assay, therefore was not implemented.

## RESULTS AND DISCUSSION

The patient derived organoids used in this study exhibited noticeable morphological differences in size, shape and aggregation properties during expansion, as shown and described in Figure 1. This brings a challenge for confocal fluorescence imaging as the aforementioned parameters influence re-distribution of the organoids during assay plating, culture and washing steps after fixing. The extent of this variation was reduced by controlling the size of the organoids to between 20 μm and 70 μm in diameter before seeding. However, optimisation was still required of not only the assay culture conditions and time course, but also the imaging settings to develop a global work flow to facilitate scaling-up and screening. Our work reported here, therefore, was focused on finding universal conditions such that the fixed endpoint parameters were covering detection of the morphological, textural and intensity properties of all 4 ovarian cancer organoid lines with the same 3D imaging and analysis settings. It is also worth noting that although culture parameters and dosing regimen may be different, it is possible to utilise this assay once organoid size and morphology is optimised to be somewhat consistent at the point of fixing. Further studies involving use of patient derived organoids from other tissue types and a variety of other compounds, not reported here, indicated a broad application of the protocol which can facilitate a wider comparison of targets.

Cell viability is one of the most common assays in drug screening in organoids ^19^ and compared to cell specific *in situ* fluorescent dye based viability measurements, for example, plate- reader based luminescence measurements reporting on cell health are more often preferred as they generally facilitate more robust data, when only well based measurements are of interest. Organoid lines were tested for viability in replicates in parallel to the imaging assay ensuring DNA damage response measurements were done mostly in viable cells. Figure 4 shows the viability of the 4 organoid lines that were exposed to the maximum concentration of Etoposide (30 μM) over the dosing period of 2 days in parallel to the imaging assay. Although variable, viability but was maintained above 72% for all the four models demonstrating sufficiently good health of the cells within the organoids after 2 days treatment for DNA damage response analysis.

**Figure 4.**
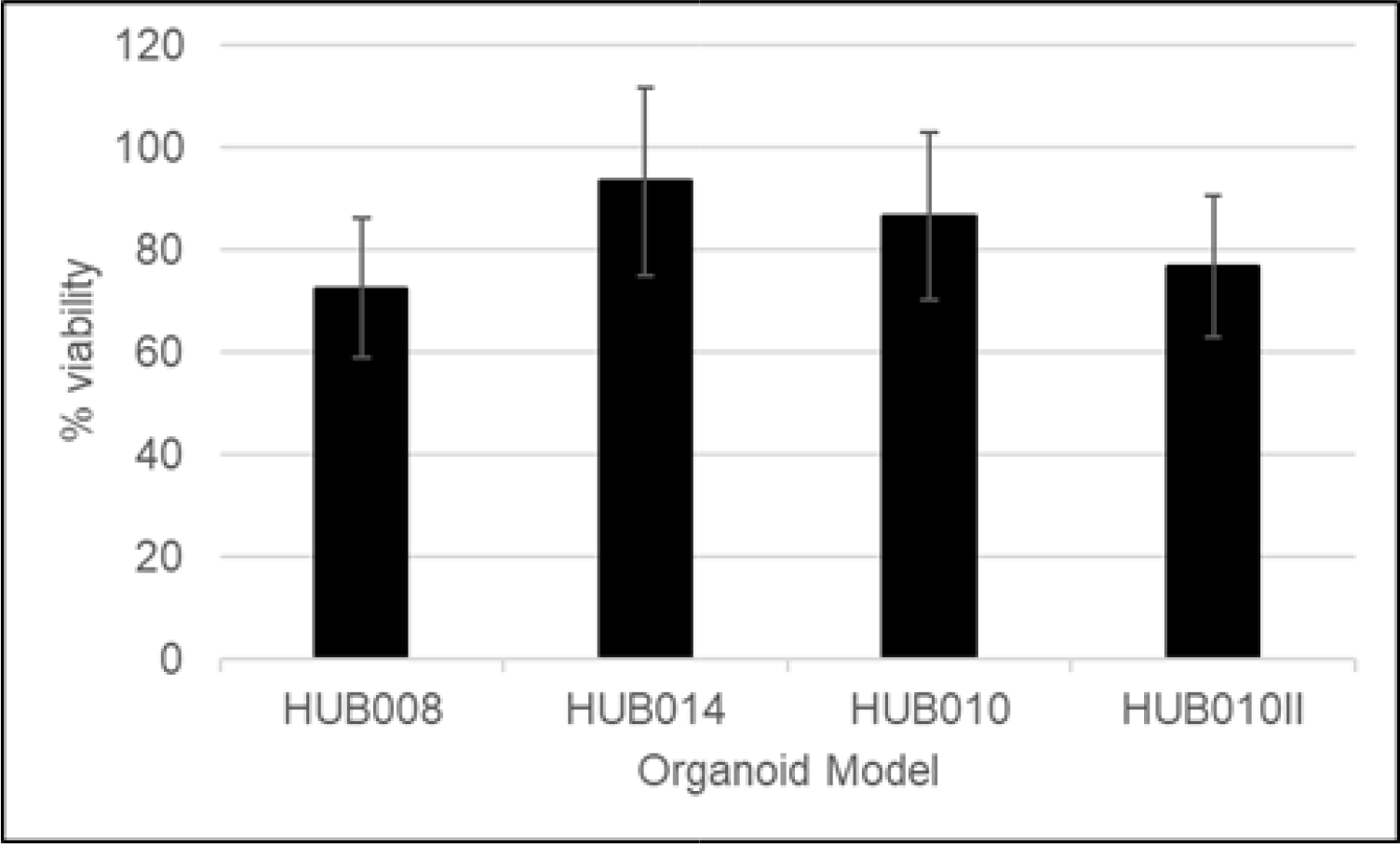
Percent viability of organoid lines following 30 μM Etoposide treatment for 2 days. Error bars demonstrate standard deviation between repeated 32 replicate measurements.

Figure 5 shows the normalised Etoposide DNA damage dose response curves fitted using linear regression option in High Content Profiler package, per organoid line with standard deviation. A good correlation between increasing Etoposide concentration and increasing DNA damage detection can be seen from different plates for HUB008, HUB010 and HUB010 II. There is relatively less response in HUB014 line, particularly at the lower to mid concentrations tested below 3.75 μM. This can also be observed in individual graphs plotted per plate for each organoid line in Supplementary Figure S3. Overall, poor curve fits for HUB014 were seen. This finding could be explained by the fact that the HUB014 organoid line is derived from a recurrent tumour following standard of care treatment and could therefore indicate establishment of drug resistance. Dose response curves for each individual organoid line are plotted in Supplementary Figure S4 for alternative visualisation.

**Figure 5.**
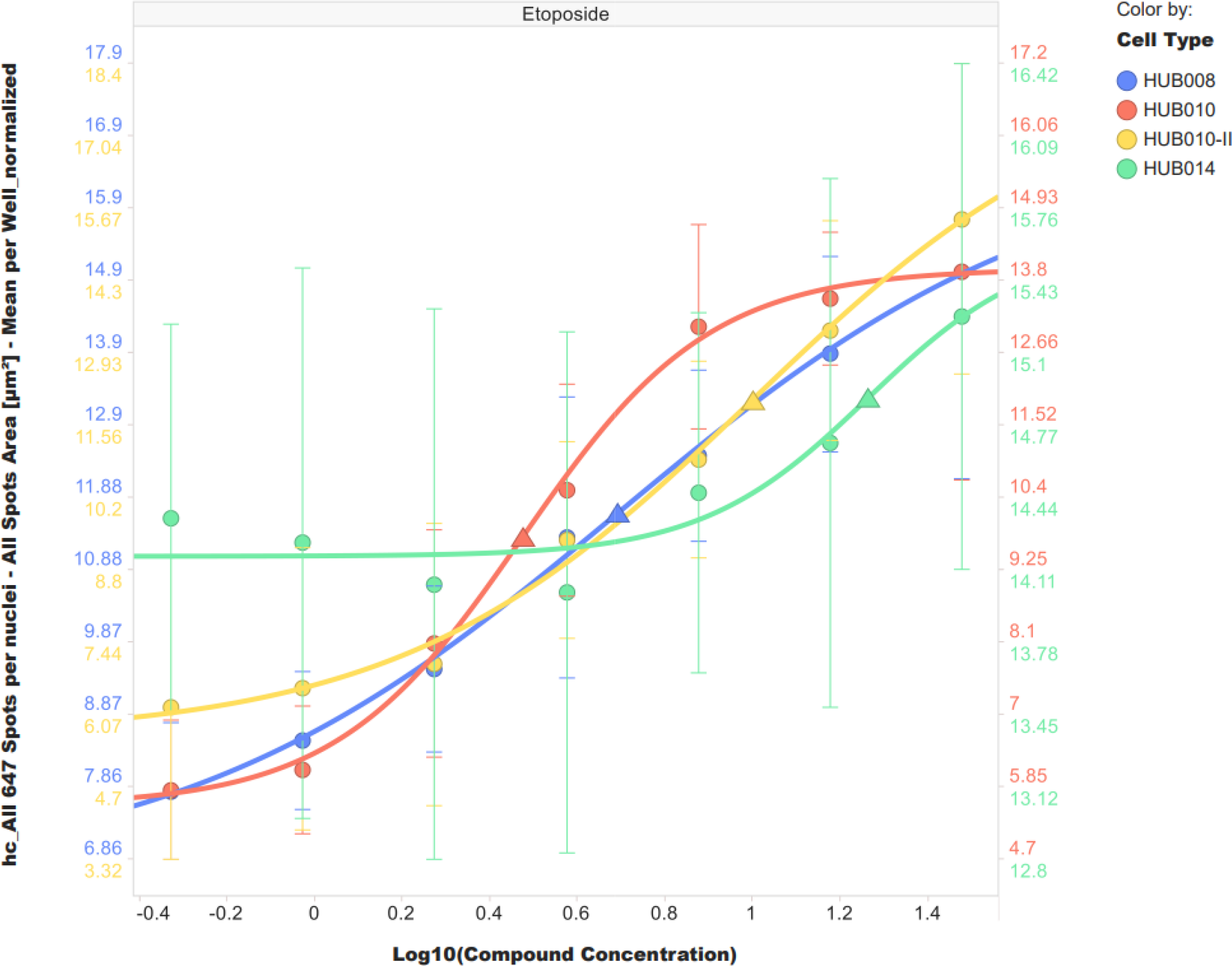
Normalised Etoposide DNA damage dose response curves per organoid line as fitted for the parameter ‘Total nuclear γH2AX spot area- mean per well’ using logistic regression model in high content profiler application. Triangle on each curve show corresponding estimated inflection point.

## CONCLUSIONS

A subnuclear 3D imaging assay that is unbiased between 4 morphologically different ovarian cancer organoid lines was outlined in the form of a method aiming broad application for wider use. Although the data reported herein is tissue specific, the 3D imaging assay is transferable between tissue types if cell density, dosing regimen and organoid size can be relatively controlled as demonstrated. A fixing, staining, and clearing protocol that is applicable to all multi-well plate based immunofluorescence organoid or spheroid assays is validated for DNA damage response. However, it should be further tested for other applications as antibody penetration and binding properties may vary and compromise quantification at scale for other markers, in particular for cytoplasmic proteins which ought to facilitate less SNR when images are projected prior to cell segmentation. The four organoid lines studied here shown expected response to Etoposide, and HUB014 has showed no quantifiable response possibly explained by its clinical origin. Our current work at GSK also involves exploring suitability of this protocol for screening other tissue types and validating targets for drug combinations in CRISPR edited organoids.

## ASSOCIATED CONTENT

Supporting Information (S)

## AUTHOR INFORMATION

Corresponding Author

H.K., C.S., and H.R. designed, performed, and analysed the experimental work and wrote the manuscript.

L.M. and E.E.E. contributed to design and writing of the manuscript.

## ACKNOWLEGEMENTS

The authors would like to thank Katja Remlinger for statistics discussions.

## DECLERATION OF CONFLICTING INTERESTS

The authors declared no conflict of interest with respect to the research, authorship, and/or publication of this article. The authors declare no competing financial interest.

## ETHICS STATEMENT

The human biological samples were sourced ethically, and their research use was in accord with the terms of the informed consents under an IRB/EC approved protocol.

