## Supplementary Figures and Tables for "A Scalable 3D High-Content Imaging Protocol for Measuring a Drug Induced DNA Damage Response Using Immunofluorescent Sub-nuclear γH2AX Spots in Patient Derived Ovarian Cancer Organoids"

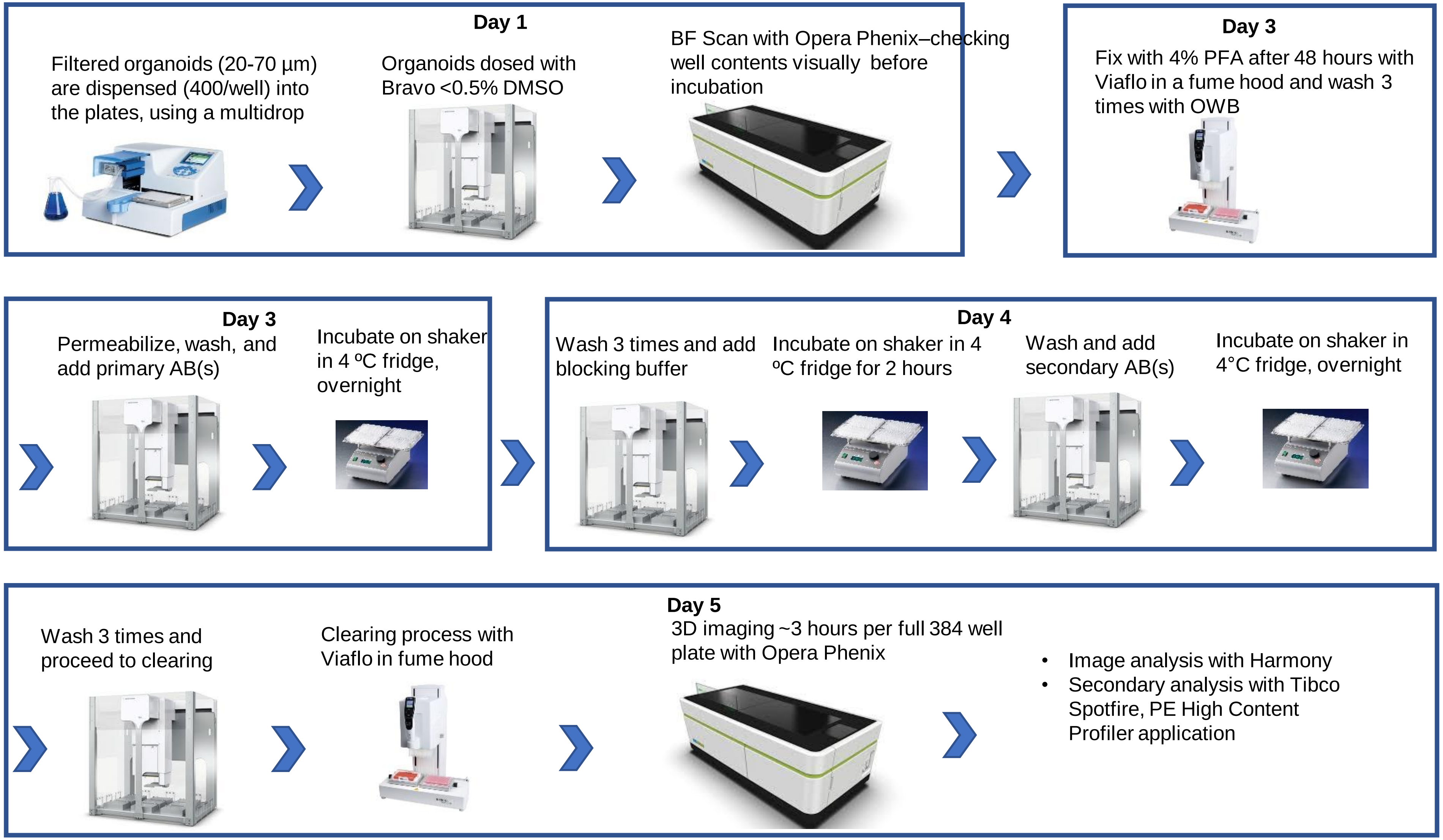


Figure S1. Organoid fixing, staining and clearing protocol workflow

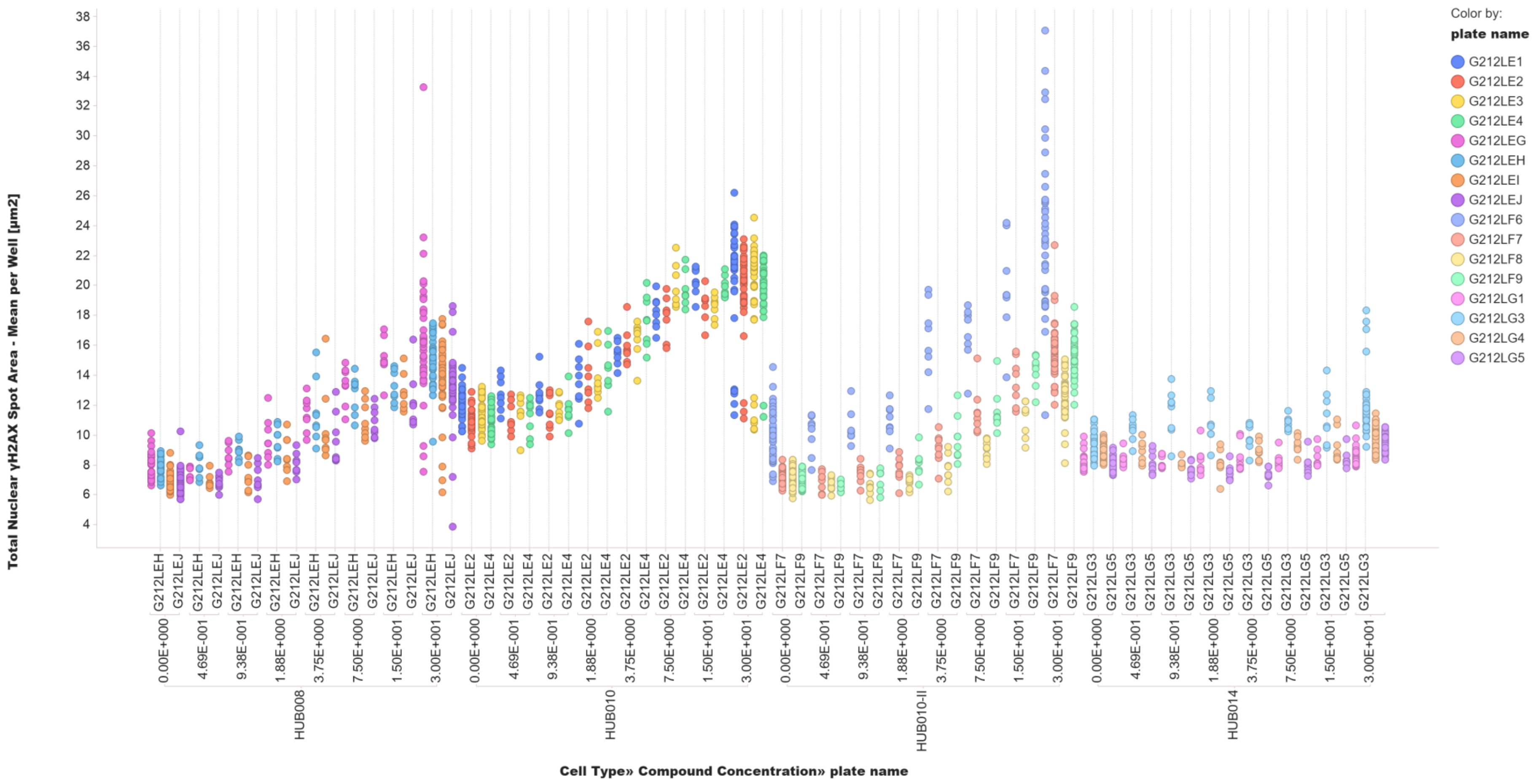


Figure S2. Raw dose response data (Total Nuclear γH2AX Spot Area- Mean per Well versus Etoposide concentration) plotted per organoid line per plate. Plate barcodes with corresponding data point colours are given in the legend (plate names are cropped on x-axis).

| # | Consumable Name | Supplier | Product Code |
|---|---|---|---|
| 1 | Methanol | Merck | 34860-100ML-R |
| 2 | CytoVista™ Invitrogen™ 3D Cell Culture Clearing/Staining Kit | Invitrogen™ | V11325 |
| 3 | BSA | Sigma Aldrich | A9647 |
| 4 | Triton X-100 | Sigma Aldrich | T8787-100ml |
| 5 | Tween 20 | Sigma Aldrich | P1379 |
| 6 | Alexa Fluor Plus 647 Goat Anti-mouse primary antibody | Thermo Fisher Scientific | A32728 |
| 7 | Phospho-Histone H2A.X (Ser139) Antibody | Thermo Fisher Scientific | MA5-27753 |
| 8 | Dapi Solution (1mg/ml) | Thermo Fisher Scientific | 62248 |
| 9 | 384 Cell Carrier Ultra ULA plates | Perkin Elmer | 6057302 |
| 10 | Sterile water | Sigma-Aldrich | W3500 |
| 11 | Nalgene™ Rapid-Flow™ Sterile Single Use Vacuum Filter Units 500 ml | Thermo Fisher Scientific | 10229090 |
| 12 | Dulbecco's Phosphate-Buffered Saline (DPBS) | Gibco | 14190-136 |
| 13 | Dispase | Gibco | 17105-041 |
| 14 | pluriStrainer 20 µm | pluriSelect | 43-50020-01 |
| 15 | CellTitre-Glo 3D | Promega | G9682 |
| 16 | 6 well suspension plate | Greiner | 657185 |
| 17 | pluriStrainer 70 µm | pluriSelect | 43-50070-01 |
| 18 | 384 well cell culture microplate, small volume, white | Greiner | 784080 |
| 19 | Nalgene Disposable Robotic Reservoir | Thermo Fisher Scientific | 1200-1301 |
| 20 | 384 Perkin Elmer Cell Carrier Ultra ULA plate | Perkin Elmer | 6057302 |
| 21 | Growth Factor Reduced Matrigel | Corning | 356231 |
| 22 | TrypLE Express Enzyme (100 mL) | Gibco | 12604013 |

Table S1. List of consumables

A)

| Organoid Line | Plate Barcode | Z-prime | Signal to Noise Ratio |
|---|---|---|---|
| HUB 008 | G212LEG | 0.2 | 3.6 |
| | G212LEH | 0.1 | 3.4 |
| | G212LEI | 0.3 | 4 |
| | G212LEJ | 0.2 | 3.5 |
| HUB 010 | G212LE1 | 0.4 | 4.7 |
| | G212LE2 | 0.1 | 3.3 |
| | G212LE3 | 0.5 | 5.9 |
| | G212LE4 | 0.5 | 5.5 |
| HUB 010 II | G212LF6 | 0.2 | 4 |
| | G212LF7 | 0.1 | 3.4 |
| | G212LF8 | 0.2 | 3.8 |
| | G212LF9 | 0.3 | 4.7 |

B)

| Top 7 Parameters (mean per well), Z-prime >0 |
|---|
| Total nuclear γH2AX spot area |
| Intensity per γH2AX spot [%CV]-mean per nucleus |
| Contrast per γH2AX spot-mean per nucleus |
| Roundness per γH2AX spot-mean per nucleus |
| γH2AX spot Haralick Homogeneity (1.19 µm) |
| Nuclei roundness |
| Nuclei area |

Table S2. Z-prime and SNR scores calculated for each plate for the organoid lines which responded to Etoposide (A), top 7 intensity and morphology features between these 3 organoid lines (B).

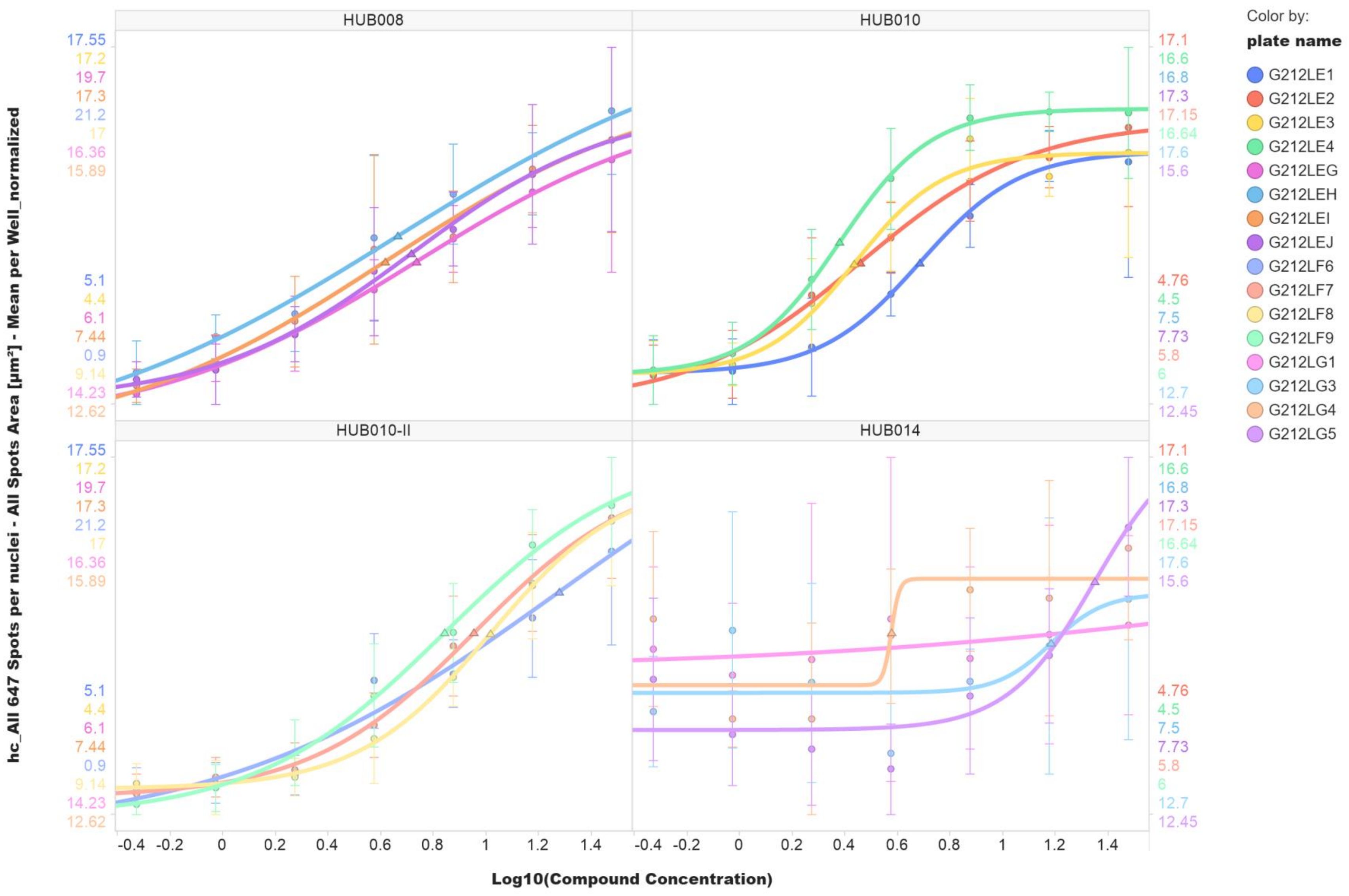


Figure S3. Etoposide dose response curves for individual plates per organoid line as fitted for the parameter ‘total nuclear γH2AX spot area- mean per well’ using the default option (linear regression) in High Content Profiler application.

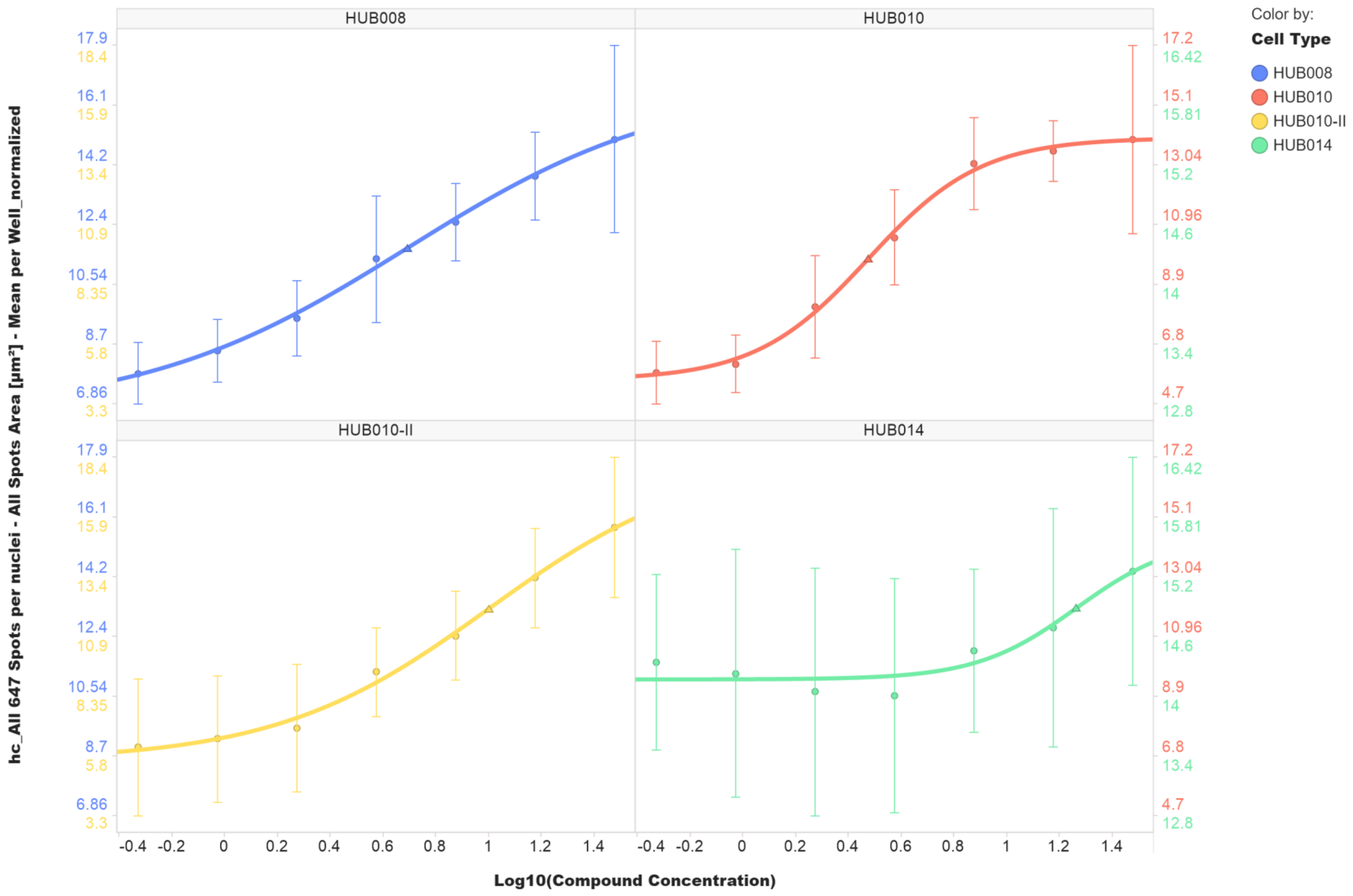


Figure S4. Etoposide dose response curves shown in Figure 5, plotted separately for each line, for alternative visualisation.
